# Maximal Efficacy of Alternative Splicing is Ensured by Balanced Efficiency of U1 and U2AF

**DOI:** 10.1101/2025.04.24.650448

**Authors:** Jianfei Hu, Daniel C. Douek

## Abstract

Alternative splicing (AS) significantly increases the diversity of gene function by enabling a single gene to produce multiple protein isoforms. U1 and U2AF are proteins responsible for the recognition and splicing of an intron’s donor and acceptor sites, respectively. Here, we used RNA-seq to explore these fundamental splicing events to separately calculate the splicing efficiency of U1 and U2AF. The expression levels of splice sites were directly calculated by the number of uniquely mapped reads spanning the splice junction. We found that the scaled expression levels of donor and acceptor sites both follow a type III extreme value Weibull distribution with the same shape parameter of 0.14. These observations significantly extend our previous findings by revealing that AS follows a Weibull distribution at the levels of both the fundamental splicing event and the mature transcript isoform. A simple transform of the shape parameter *a*, 1/(1+*a*), ranges between 0 and 1, and positively correlates with the dominance of the major splicing product and therefore represents an index of the splicing machinery efficiency. Importantly, the equal splicing efficiency of U1 and U2AF ensure that their combined efficacy reaches a maximum.

## Introduction

Alternative splicing (AS) of genes is a critical process for eukaryotic organisms. It enables a single gene to generate multiple different transcript isoforms that further encode multiple different proteins with potentially different functions; thus, the total number of distinct proteins that a human cell can produce is far higher than the total number of genes in the human genome (*1*). Many studies have identified cis-acting elements and trans-acting factors, as well as specific accessory proteins involved in AS (*2-5*). The details of some of the molecular events have also been elucidated. The U1 protein binds to the 5’ end of an intron, recognizing the donor sites, while U2AF (the complex of U2AF^65^ and U2AF^35^) binds to the 3’ end of an intron to recognize the acceptor sites. This binding of U1 and U2AF is ATP-independent, weak and reversible, and becomes stable only after the ATP-dependent binding of U2 snRNP. U1 is then released and other accessory proteins such as U2, U4- 6 snRNP and SR are recruited to form serial A/B/C/P complexes that complete intron splicing (*6- 9*). Stronger binding of U1 and U2AF to the splice sites allows more time for the formation of stable complexes, thus increasing the likelihood that the corresponding intron will be spliced (*10*). Mathematically, this process represents a stochastic minimization process in which U1 and U2AF dynamically search their global or local minimal potential energy sites on pre-mRNA. This minimization process usually leads to an extreme value distribution as its outcome (*11-14*). RNA-seq offers both single-base resolution for annotation and ‘digital’ expression level measurement of transcript isoforms at the genome scale, thus enabling accurate mathematical analysis of AS products to derive the physical characteristics of AS mechanisms (*10*). Previously, we have proposed a mathematical model to describe AS which shows that the scaled expression levels of transcript isoforms from different genes all follow the same type III extreme value distribution — the Weibull distribution given in formula 1 — that can further describe the frequency distribution of transcript isoforms as well as many previously unexplained observations such as that genes tend to express all their isoforms simultaneously but at different levels (*14*).

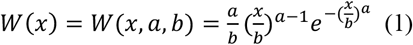

*W(x)* is the probability of transcript isoform with expression level x; *b* is the scale parameter which will change with the expression level of gene; and *a* is the shape parameter which is specific for the AS mechanism. Simulation analysis also revealed an interesting characteristic of the Weibull distribution which ensures that the major transcript isoforms of a gene have a low probability of dropout at the single cell level (*15*).

The recognition and splicing of single 5’ donor and 3’ acceptor sites are the most basic AS events. The expression level of a splicing event can be defined as the number of uniquely mapped reads spanning the splice junction. Previous analyses used data generated from Illumina next-generation sequencing (NGS) libraries which are typically of insufficient read length (100 - 300bp) to cover an entire transcript isoform. Thus, statistical inference such as an Expectation Maximization (EM) algorithm must be used to assemble transcript isoforms and calculate their expression levels indirectly (*16-19*). However, these read lengths are quite sufficient to detect novel basic splicing events and directly calculate their expression levels, which enables us to analyze the physical characteristics of AS events at a fundamental level. Five major types of AS events have been identified: exon skipping, mutually exclusive exons, alternative 5’ donor sites, alternative 3’ acceptor sites and intron retention (*1, 4, 20, 21*). Of these, alternative 5’ donor sites and alternative 3’ acceptor sites are of particular interest as they enable us to analyze the separate contribution of U1 and U2AF in the AS process.

## Results

### Alternative donor and acceptor sites are common in AS

Let us suppose that D1 is a donor site with four alternative acceptor sites: A1, A2, A3 and A4 (Fig 1A). The expression level of the D1_A1 splicing event is determined by the binding strength of U1 to D1 and U2AF to A1 as well as the overall expression level of the gene in which these splice sites are located. The latter factor is identical for all splicing events in the gene and thus can be disregarded here. The expression level of D1_A2 is thus determined by the binding strength of U1 to D1 and U2AF to A2, and the same principle applies to the D1_A3 and D1_A4 splicing events. The binding strength of U1 to D1 is same for all four splicing events, and so any differences in their relative expression level is uniquely determined by the different binding strength of U2AF to A1, A2, A3 and A4. The analysis of the relative expression level of these four splicing events enables us to probe the physical characteristics of U2AF binding. Similarly, splicing events with alternative 5’ donor sites share the same 3’ acceptor sites; thus, the relative expression levels of these splicing events exclude the influence of U2AF binding and is solely influenced by U1 binding of the different 5’ donor sites (Fig 1B). Thus, we analyzed the expression levels of alternative 5’ donor and 3’ acceptor sites by RNA-seq to understand the physical characteristics of U1 and U2AF binding, and to determine whether the Weibull frequency distribution also applies to the most fundamental level of AS.

**Fig 1.**
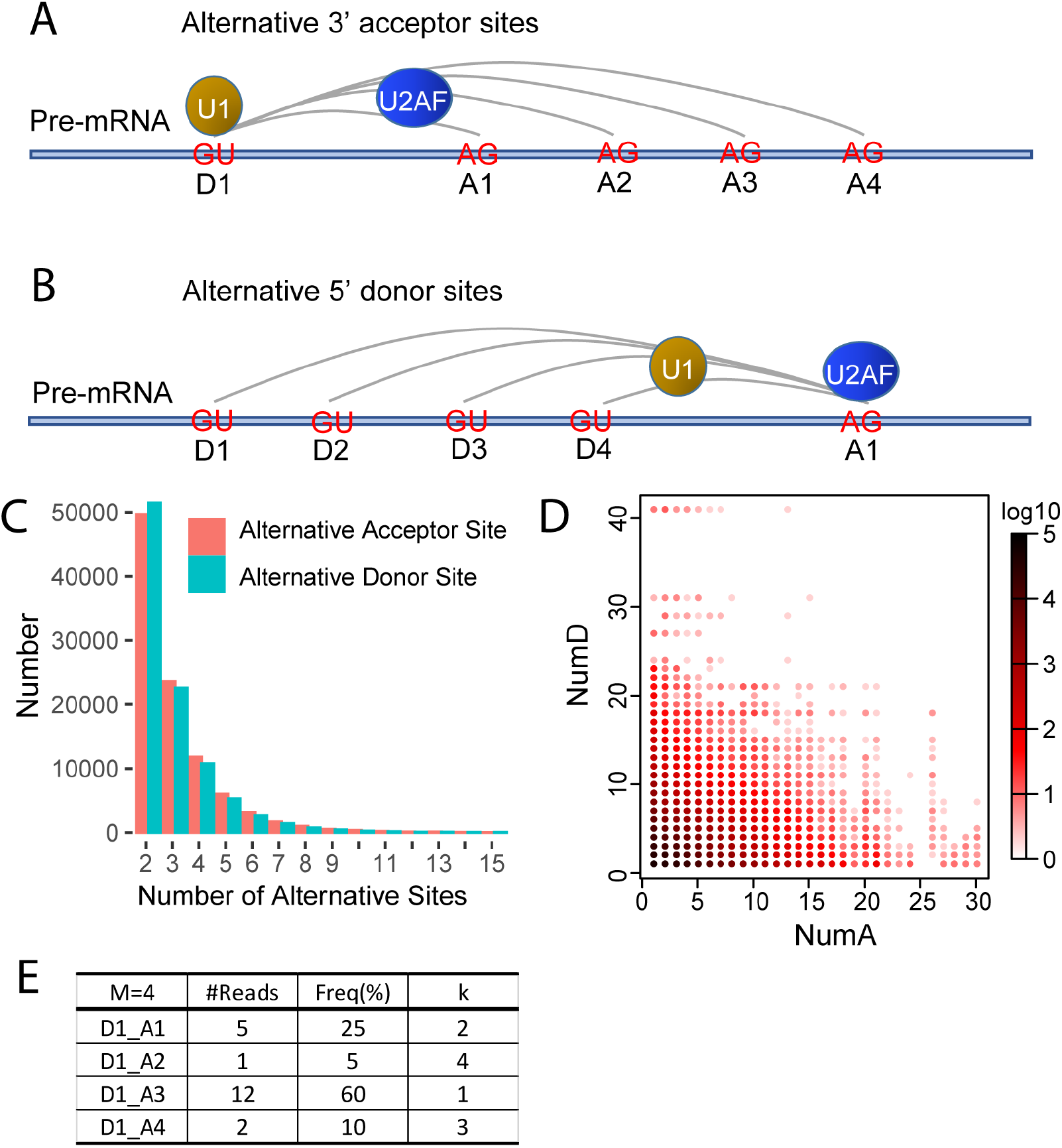
Alternative donor and acceptor sites. (A) Schematic figure of alternative 3’ acceptor sites. U1 protein binds donor site D1; U2AF protein may bind to acceptor site A1, A2, A3 or A4. (B) Schematic figure of alternative 5’ donor sites. U1 protein may bind to donor site D1, D2, D3 or D4; U2AF protein binds to acceptor site A1. (C) Histogram of the number of alternative donor/acceptor sites. (D) Heatmap of the number of alternative donor/acceptor sites. Supposing D_A is a splice junction, donor site D corresponds to numA different alternative acceptor sites, acceptor site A corresponds to numD different alternative donor sites and the coordinates in the graph will be (numA, numD). (E) Schematic diagram of calculation of frequencies of alternative acceptor site. M is the total number alternative acceptor sites associated with this donor site. k is the rank of the acceptor sites.

Bulk RNA-seq data of highly purified human CD4 T cell subsets deposited in public GEO database (GSE80016) were used in our analysis, which included 72 samples from nine different donors (*14*). A total of 471,325 high quality splice junctions with gene strand information were detected in these 72 samples. 99% of the splice junctions had maximum spliced alignment overhangs greater than 20bp. Each splice junction was detected in at least three independent samples (see Methods for details). 96,044 donor sites had more than one corresponding acceptor site (i.e.: alternative acceptor sites) and 98,372 acceptor sites had more than one corresponding donor sites (i.e.: alternative donor sites). The number of donor sites with 2, 5, 10 and 15 alternative acceptor sites were 49,671, 6029, 346 and 36, respectively. The number of acceptor sites with 2, 5, 10 and 15 alternative donor sites were 51,444, 5279, 261 and 37, respectively (Fig 1C). Of the 471,325 high quality splice junctions detected, only 62,473 (13.3%) had no alternative splice sites at either end. The remaining 86.7% of splice junctions either had a donor site with alternative acceptor sites, an acceptor site with alternative donor sites, or alternative splice sites at both ends (Fig 1D). These results indicated that alternative donor and acceptor sites are common. In the following analyses, the maximum number of alternative donor and acceptor sites were set at 15 as few donor and acceptor sites have more than 15 alternative splice sites.

We defined the frequency of an alternative acceptor site as the ratio of its expression level relative to the total expression level of all alternative acceptor sites sharing the same donor site. Thus, for a donor site with M different alternative acceptor sites where each acceptor site has the rank *k* in the hierarchy of expression level, we use *f*(*k, M*) to represent the frequency of the *k*_th_ dominant acceptor sites. As 1≤*k*≤*M*, so *f(1, M)* ≥ *f(2, M)* ≥ … ≥ *f(M, M). f(k, M)* was entirely stochastic, differed among donor sites and changed with the expression level of donor sites. For example, *f(1,2)*, the frequency of the most dominant acceptor site of a donor site with two alternative acceptor sites, varied between 50% and 100%. *f(2,2)*, the frequency of the second most dominant acceptor site, varied between 0 and 50%. The definition of *f(k, M)* for alternative donor sites is the same as for alternative acceptor sites (Fig 1E).

### The scaled expression levels of alternative donor and acceptors sites follow the same Weibull distribution

To analyze the probability density distribution of the scaled expression level of alternative acceptor sites from different genes, we first normalized them using a simple scaling factor comprising the mean expression level of all alternative acceptor sites sharing the same donor sites. The red and blue curves in Fig 2A show the density plots of normalized expression levels for alternative acceptor and donor sites, respectively. It is immediately apparent that even though the two curves contain many peaks, they nevertheless almost exactly overlap both with each other and with the black curve representing the simulated distribution (described below). These peaks were introduced as a consequence of scaling of the expression levels and their positions correspond to the different integer value of *M*, the number of alternative acceptor and donor sites.

**Fig 2.**
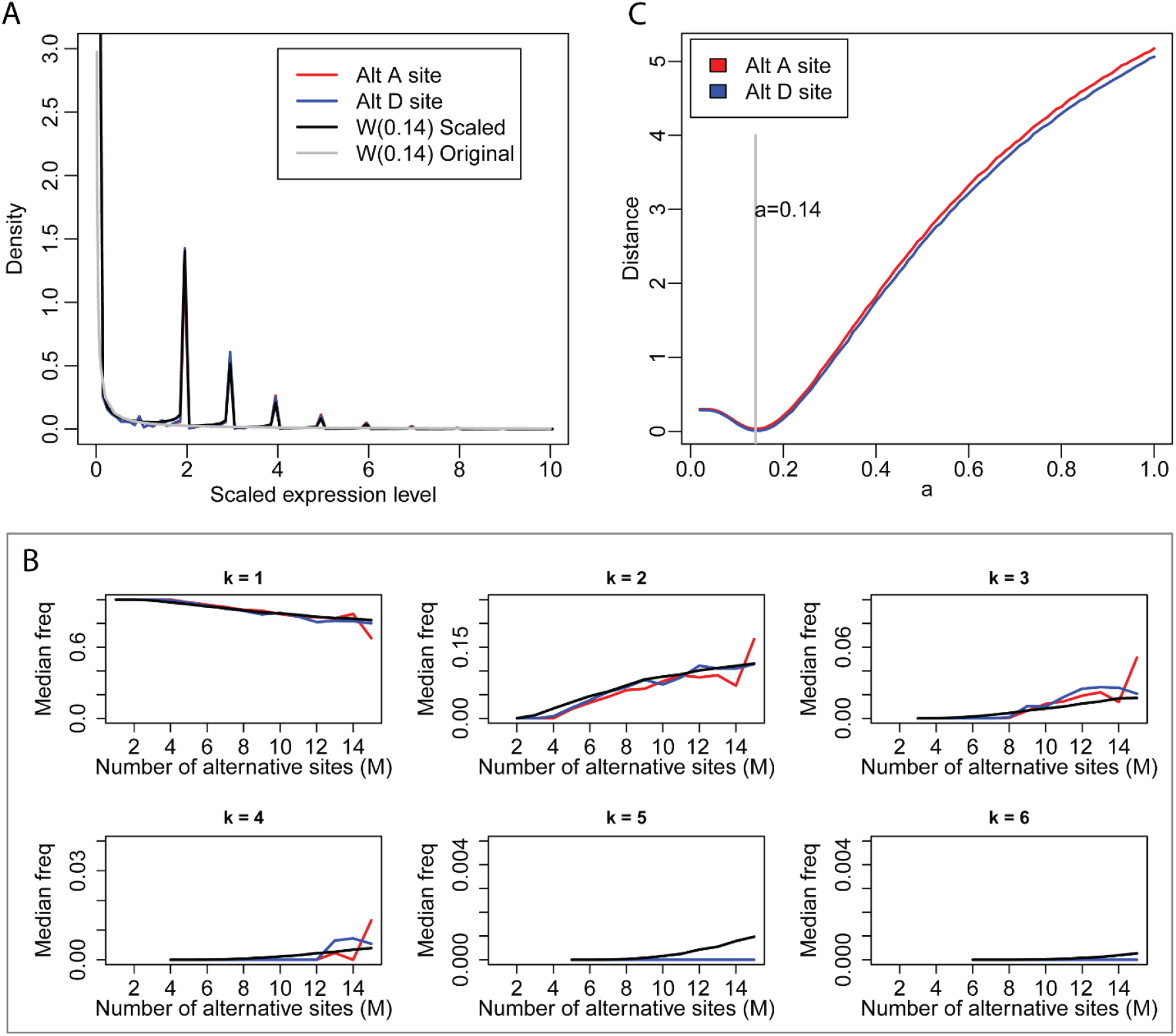
The distribution of scaled expression level and frequency of alternative acceptor and donor sites. (A). The distribution of scaled expression level. (B). The distribution of median frequency of alternative acceptor and donor sites with fixed *k* and increasing *M* from RNA-seq data. *k* is the rank of alternative splice sites. *M* is total number of alternative splice sites. (C). The distribution of the Euclidian distance relative to different *a* for all *mf(k,M)* in Fig 2B between experimental data and simulated data from Weibull distribution. The distance reaches the minimum when *a*=0.14 for both alternative acceptor sites and donor sites. Red line represents alternative acceptor sites. Blue line represents alternative donor sites. Black line represents simulated distribution from *W*(0.14). Gray line is density curve of *W*(0.14).

To analyze the frequency distribution of alternative acceptor sites, we grouped donor sites according to the number of associated alternative acceptor sites, beginning with those associated with two through *M* alternative acceptor sites. The variation of frequency with dominancy rank *k* and *M* is shown as boxplots in Fig S1. The frequency of the most dominant acceptor site (*k*=1) decreases slowly with M, while the frequencies of other acceptor sites (*k*=2, 3, 4) increase slowly with M. Similar analyses performed on alternative donor sites produced the same result (Fig S2). To compare the change in frequency for *k* and *M* for alternative acceptor and donor sites simultaneously, we extracted the median frequency *mf(k,M)* for both and plotted them on the same graph (Fig 2B). The two groups of curves corresponding to alternative acceptor and donor sites almost completely overlapped with each other.

Because of the scaling effects and peaks in the curves, we could not use curve fitting to calculate the shape parameter of the Weibull distribution. Thus, we used a Monte Carlo simulation to calculate the accurate value of shape parameter *a* (*14*), with the scale parameter *b* set at 1 as it has no influence on the calculation. We randomly selected a number in the range (0, 1) as the value for *a* and performed the following computation: for donor sites with *M* alternative acceptor sites, we randomly extracted *M* numbers from the Weibull distribution *W(a,1)* as the expression level of the *M* simulated acceptor sites, which could then be transformed to their frequencies. We repeated this process 10,000 times for each *M* to obtain the simulated median frequency *mf(k,M)* and then compared it with the corresponding *mf(k,M)* from our experimental data. The Euclidian distance of all *mf(k,M)* from simulated data and experimental data in Fig 2B was calculated and we found that it reached the minimal value when *a* was 0.14 (red curve in Fig 2C). We then performed the same analysis for alternative donor sites and found that the Euclidian distance also reached the minimal value when *a* was 0.14. The black curve in Fig 2B shows that the simulated *mf(k,M)* from *W*(0.14) overlaps well with the red and blue curves from the experimental data when *k* is 1, 2 or 3. When *k* is larger than 4, *mf(k,M)* is very small because sequencing depth is not sufficient to detect rare splice sites; thus, *mf(k,M)* from the experimental data is always 0, while the simulation from W(0.14) may still produce theoretical values.

Using the shape parameter *a*=0.14, the experimental data (red and blue curves) in Fig 2A may be exactly reproduced by the Monte Carlo simulation. For donor sites with M alternative acceptor sites, we randomly extracted M numbers from the Weibull distribution W(0.14) as the expression level of the M simulated acceptor sites, we then normalized these M numbers by their mean value to obtain their scaled expression levels. We then combined these scaled expression levels for different M to produce the density plot. The simulated density plot (black curve in Fig 2A) almost exactly matched with that of the experimental data for the alternative acceptor and donor sites (red and blue curves in Fig 2A).

Thus, we showed that the scaled expression level of alternative donor and acceptor sites also follows a Weibull distribution. Taken together, these new findings and our previous observations (*14*) demonstrate that the Weibull distribution applies at the level of both transcript isoform and basic splicing events. Nonetheless, the term *W(x)* in formula (1) has different connotations at these two levels. For alternative donor and acceptor sites, *W(x)* is the probability of a donor site or an acceptor site with expression level x; *b* is the scale parameter, which will change with the expression level of related donor sites or acceptor sites; and *a* is the shape parameter, which is specific for the binding affinity of the U1 and U2AF protein.

### The frequency distribution of alternative donor and acceptor sites

Our model was able not only to explain the median frequency *mf(k,M)* of alternative donor and acceptor sites, but also to provide the statistical distribution of each *f(k,M)*. Repeating the previous Monte Carlo simulation with *a*=0.14, we obtained the frequency distribution of alternative splice acceptor sites and alternative splice donor sites for different groups (different *M*) (Fig 3) and each *f(k,M)* (Fig 4) from the simulated data. We used Kullback-Leibler divergence (KLd) to evaluate the difference between two distributions, which represents the amount of information lost when we used the simulated frequency distribution from our Weibull model to represent the frequency distribution from the experimental data. We found that for most frequency distributions analyzed, the amount of information lost is less than 0.1 (mean=0.081, median=0.088). This shows that the frequency distribution from the simulated data (black curve) is highly consistent with that from the experimental data (red curve and blue curve), although the shape and range of the distribution change with *k* and *M*. Thus, although the expression levels of alternative acceptor sites change with a particular donor site and the expression level of the related gene, their overall frequency distributions do not change and can be described by our model. Importantly, expression levels of alternative donor sites follow the same pattern.

**Fig 3.**
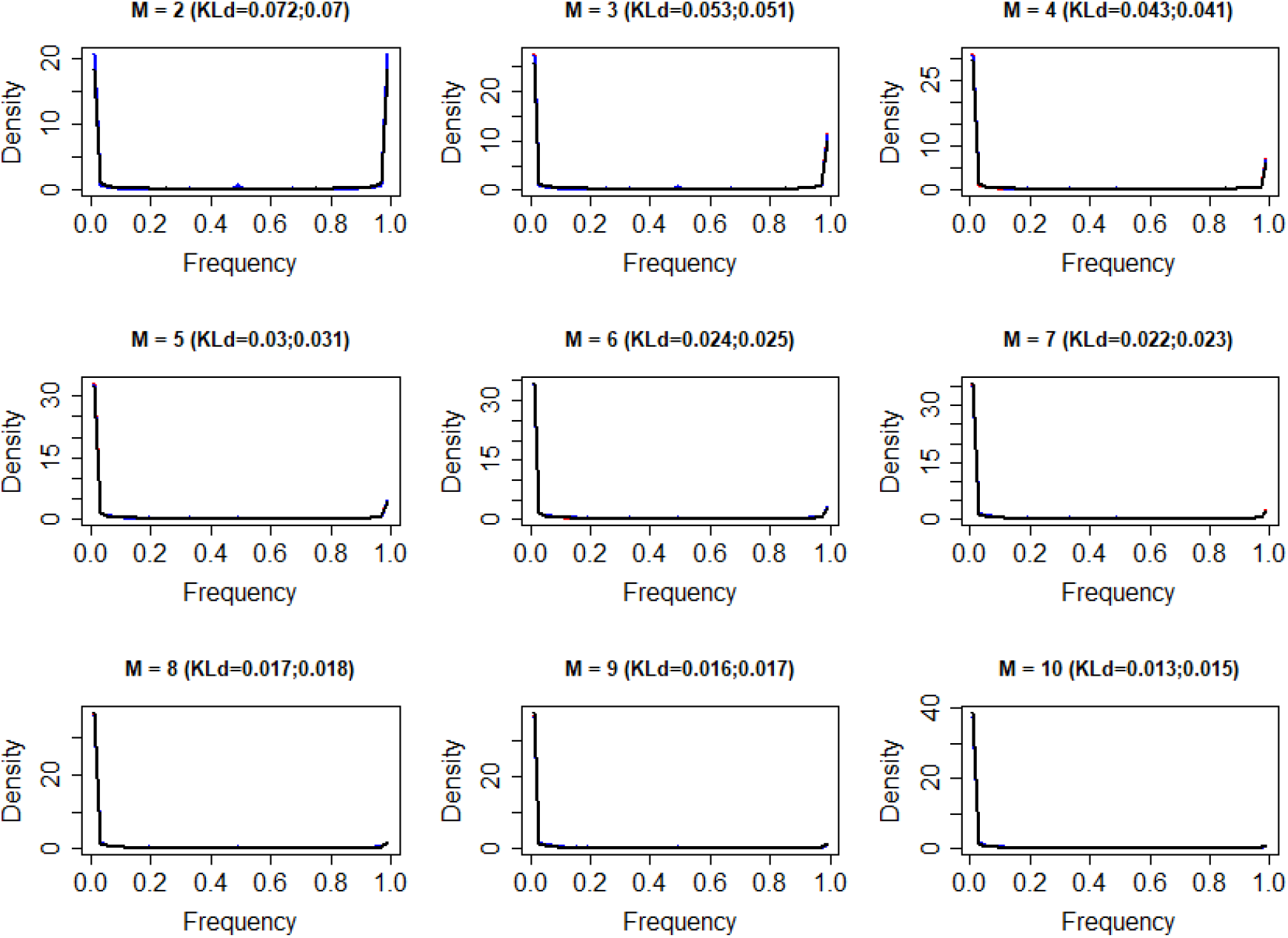
The frequency distribution of all alterative splice sites from experimental data and simulated data for M=2:10. M is the number of alternative donor and acceptor sites. Red curves represent alternative acceptor sites, blue curves represent alternative donor sites, black curves represent simulated data from *W*(0.14). KLd is the Kullback-Leibler divergence between the two distributions. The first number in parentheses is KLd between the frequency distribution of alternative acceptor sites and simulated data from *W*(0.14). The second number in parentheses is KLd between the frequency distribution of alternative donor sites and simulated data from *W*(0.14). The three curves almost exactly overlap.

**Fig 4.**
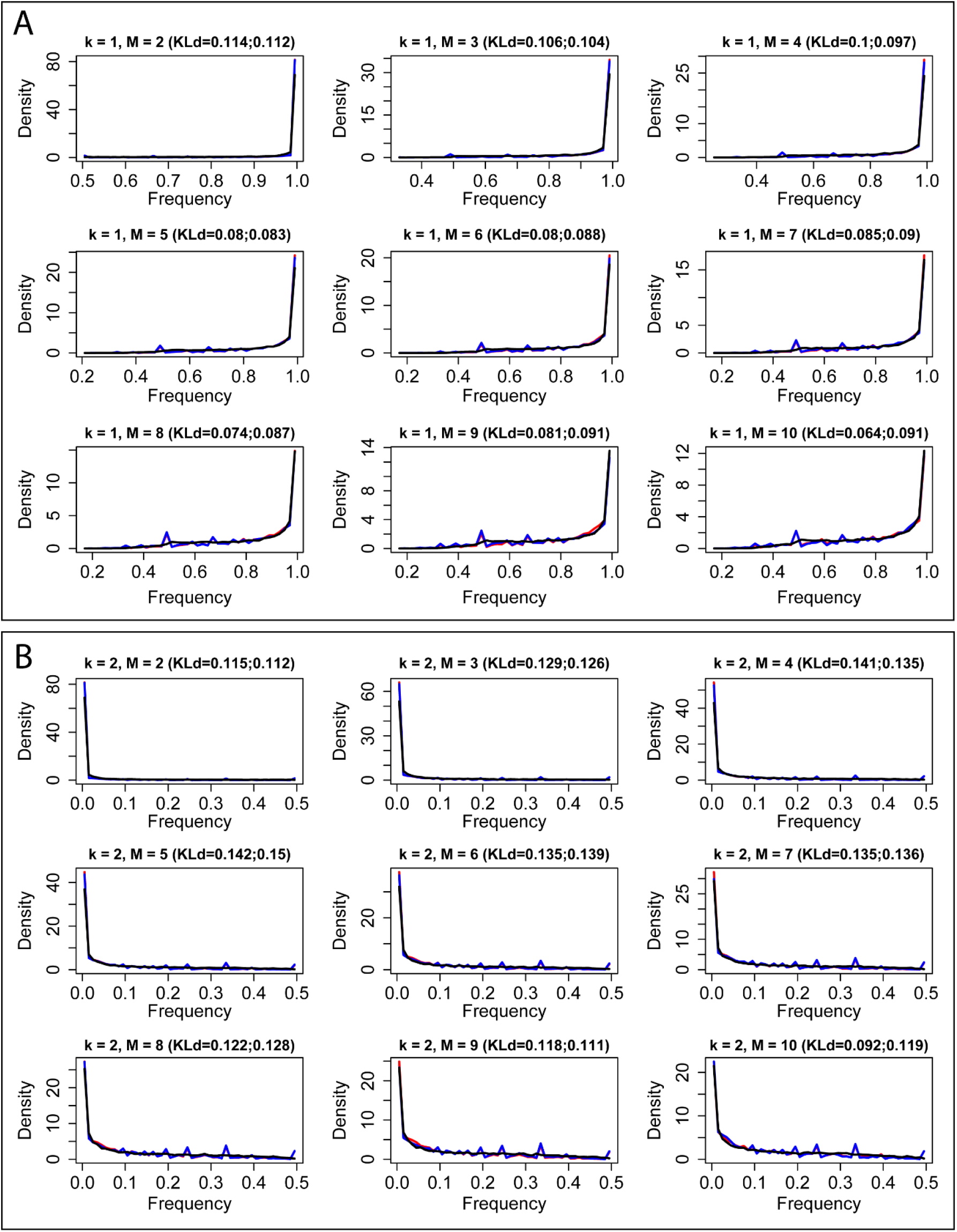
The frequency distribution of kth dominant alterative splice sites. (A) k=1. (B) k=2. M is the number of alternative donor and acceptor sites. Red curves represent alternative acceptor sites, blue curves represent alternative donor sites, black curves represent simulated data from *W*(0.14). KLd is the Kullback-Leibler divergence between two distributions. The first number in parentheses is the KLd between the frequency distribution of alternative acceptor sites and simulated data from *W*(0.14). The second number in parentheses is KLd between the frequency distribution of alternative donor sites and simulated data from *W*(0.14). The three curves almost exactly overlap.

### Verification using independent RNA-seq datasets

To confirm that the results we obtained are generally applicable and independent of the dataset used, we performed the same analysis on a colorectal cancer (CRC) RNA-seq dataset (GSE50760) from NCBI GEO which comprised 54 samples including normal colon, primary CRC and liver metastases from 18 CRC patients. We obtained similar results for all analyses performed (Fig S3- S6).

### Comparison between the number of expressed and annotated alternative donor and acceptor sites

AS is a stochastic process and thus not all annotated alternative donor and acceptor sites will be expressed in a biological sample and detected in its RNA-seq dataset. Nonetheless, our Weibull model may accurately predict the number of alternative donor and acceptor sites observed using the expression levels from the simulated Weibull distribution *W*(0.14). As different alternative donor sites have different expression levels, we could reasonably select a frequency cutoff rather than the absolute expression level to define whether an alternative donor site would theoretically be detected. The frequency cutoff used is influenced by both the sequencing depth and the number of cells in the sample, becoming lower with deeper sequencing and more cells sampled. Thus, we defined undetectable splice donor sites as those below a frequency cutoff of 0.002. The boxplots in Fig S7 and S8 show the observed result from our RNA-seq dataset and confirm good concordance between the two plots for both alternative donor sites and alternative acceptor sites.

### The physical significance of the shape parameter

AS follows a Weibull distribution both for basic splicing events and for mature transcript isoform expression, each with a different shape parameter of *W*(0.14) and *W*(0.39), respectively. To understand the physical significance of the shape parameter *a* in AS, we compared various characteristics of the Weibull distribution with different shape parameters. For convenience, we define the dominance of the major splice product as its median frequency *mf(1,M)*. First, from the probability density plot shown in Fig 5A, a smaller shape parameter *a* generates a steeper curve with a greater density around 0. Second, when *M* is fixed, the dominance of the major splice product increases as the shape parameter *a* decreases (Fig 5B). When *a* is small, such as 0.05, *mf(1,M)* is larger than 0.99 for M≥1, indicating that transcript isoforms almost entirely comprise the major one. When *a* is large, such as 1, *mf(1,M)* is less than 0.5 for M>5, indicating that the major transcript isoforms are not more dominant than other isoforms. Typically, the percent spliced in index (PSI) is the most commonly used index to indicate the efficiency of splicing a specific exon into the transcript population of gene (*22, 23*). Similarly, a simple transform of the shape parameter, 1/(1+*a*), can be defined as a novel index in AS analysis to indicate the overall efficiency of the splicing machinery in the generation of all major splicing products. Under this definition, the smaller the shape parameter *a*, the higher the splicing efficiency, which ranges from 0 to 1 and positively correlates with the dominance of the major splice product (Fig 5C and 5D). As the splicing efficiency increases from 0 to 1, the dominance of the major splice product increases from 1/*M* to 1. Depending on the context, 1/(1+*a*) can indicate the overall efficiency of: (1) U1 in splicing all major donor sites; or (2) U2AF in splicing all major acceptor sites; or (3) the entire AS machinery in splicing all major transcript isoforms. Under this definition, the efficiency of splicing at the basic level of donor and acceptor sites is 1/(1+0.14)=0.88, which is higher than the efficiency at the mature transcript isoform level of 1/(1+0.39)=0.72. Furthermore, the observation that alternative donor and acceptor events have equal shape parameters suggests that U1 and U2AF have equal efficiency in recognition of the donor and acceptor sites.

**Fig 5.**
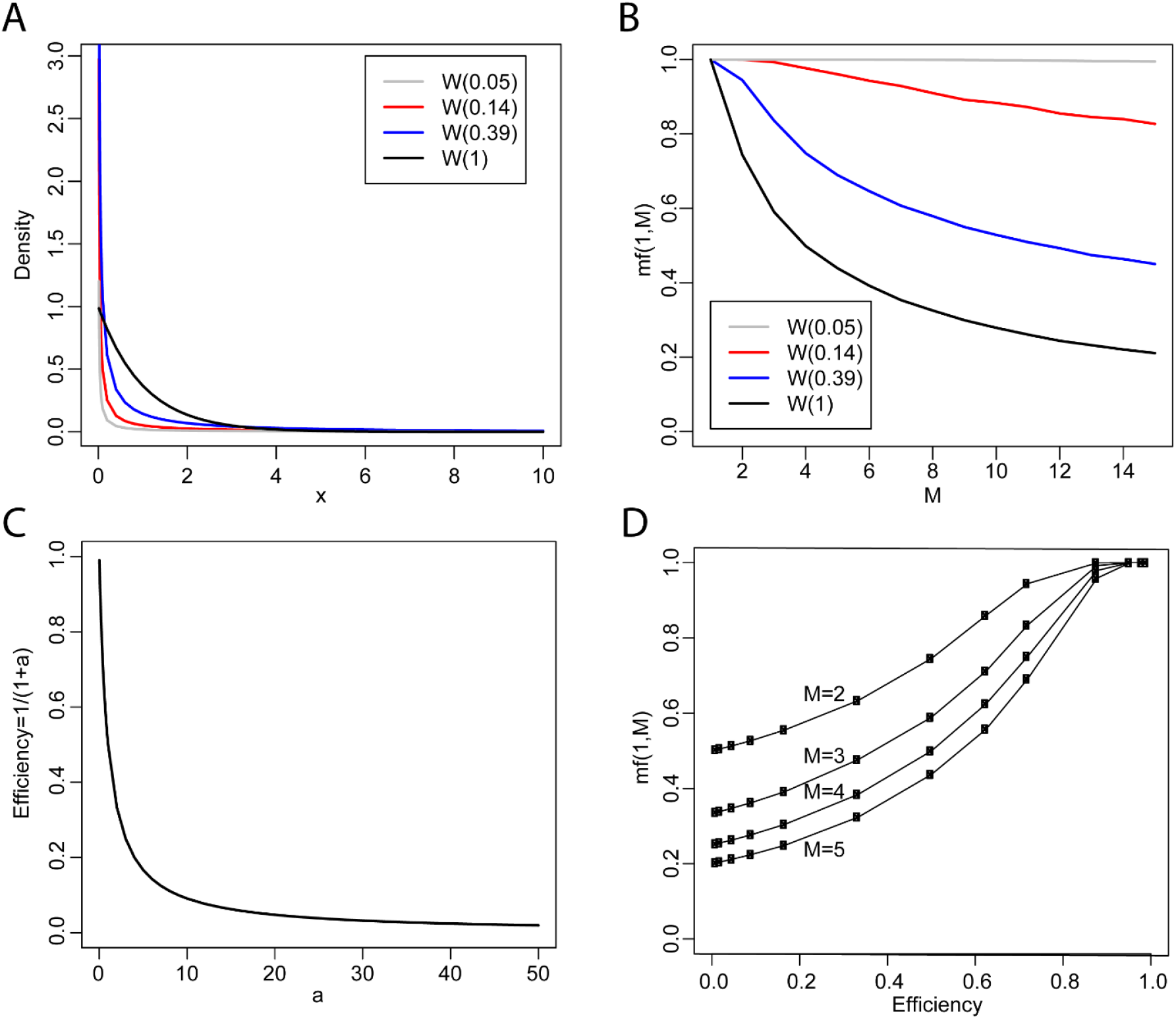
The physical significance of the shape parameters. (A). Probability density plot of Weibull distributions with different shape parameters. (B). The median frequency of the major splice product *mf(1,M)* from simulation of Weibull distribution with different shape parameter. (C). The shape parameter *a* negatively correlates with the splicing efficiency. The smaller the *a*, the higher the efficiency of splicing machinery. (D). The splicing efficiency positively correlates with the dominancy of the major splicing product. *W*(0.14) corresponds to the Weibull distribution at alternative acceptor or donor site level. *W*(0.39) corresponds to the Weibull distribution at transcript isoform level.

## Discussion

In our previous study we used bulk RNA-seq datasets to analyze the probability density distribution of scaled transcript isoform expression level and found that it follows an extreme value Weibull distribution of *W(0*.*39)*. However, because short sequencing reads (≤300bp) cannot cover the entire length of a transcript, its expression level can only be calculated indirectly by statistical inference. In the present study, we overcame this problem through analysis of the probability density distribution of the expression level of alternative donor and acceptor sites in RNA-seq datasets. Thus, the results derive directly from the mapping of sequencing reads on the human reference genome with no statistical inference or estimation in the calculation of the expression level. We found that alternative donor and acceptor sites both follow the same Weibull distribution *W*(0.14) which explains not only the distribution of their expression level but also the distribution of their frequency. Together with our previous results, we may conclude that AS follows a Weibull distribution at both the basic splicing event level and at the mature transcript isoform level. A simple transform of the shape parameter, 1/(1+*a*), represents the efficiency of splicing: the higher is the efficiency of splicing, and the more dominant is the major splicing product. When considering the molecular complexity of AS where errors may occur at each splicing event, one may hypothesize that a fully spliced transcript isoform will have experienced multiple splicing events and errors will have accumulated during this process; thus the overall efficiency at the transcript isoform level should be lower than that at the basic splicing event level — a hypothesis which is now confirmed by our results. A limitation of our analysis is that the expression level of splice sites and transcript isoforms evaluated by RNA-seq is the result of joint actions of splicing and degradation. Unfortunately, this limitation exists in all large-scale transcriptome-level gene expression analysis based on sequencing and cannot be resolved with currently available approaches. Finally, our finding that alternative donor and acceptor sites possess equal shape parameters indicates that U1 and U2AF — key proteins of the splicing machinery — possess identical efficiency in recognizing their splice sites. Supposing the splicing efficiency of U1 and U2AF are η_1_ and η_2_, the joint intron splicing efficiency of U1 and U2AF will be η_1_×η_2._ According to the inequality of arithmetic and geometric means, for two non-negative numbers, 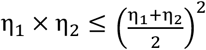, when the sum is fixed, the product of the two numbers reaches a maximum when they are equal. Thus, we conclude that equality of the individual efficiencies ensures that the combined efficacy of U1 and U2AF reaches a maximum and we show that our mathematical analysis of large-scale sequencing data can reveal physical characteristics of the complex biological process of AS.

## Dataset and Methods

### Datasets

Previously published bulk RNA-seq data of highly purified human CD4 T cells were used for the analysis (GSE80016). This dataset contained 72 samples from nine different donors. Each donor has eight samples sequenced, including four T cell subtypes (naïve, central memory, transitional memory and effector memory) and two statuses (resting and activated). These samples were sequenced in 2×100 base paired-end runs. The dataset comprised 1.27×10^9^ reads pairs in total and each sample corresponds to 1.76×10^7^ reads pairs on average (*14*).

### Bioinformatic analysis

Trimmomatic (version 0.22) was used to remove adapters and low-quality bases(*24*). The trimmed paired-end reads were mapped to reference human genome (Hg38) using STAR (version 2.7.9a)(*25*). Duplicated read pairs were removed by Picard (http://picard.sourceforge.net/, version 2.1.1). The output “SJ.out.tab” was used for AS analysis. 1,186,496 distinct splice junctions were detected in these 72 samples in total. To remove false positives, we required that each splice junction be detected in at least three independent samples. 542,390 splice junctions remained after this filtering.

If a gene is located on the positive strand of the chromosome, the left end of the splice junction is the 5’ donor site and the right end of the splice junction is the 3’ acceptor site. If a gene is located on the negative strand of the chromosome, the left end of the splice junction is the 3’ acceptor site and the right end of the splice junction is the 5’ donor site. Ensembl annotation of human genome (“Homo_sapiens.GRCh38.103.gtf”) was used for gene strand analysis. Splice junctions located in intergenic regions or in more than one gene region with conflicting gene strand information were discarded. 471,325 high-quality splice junctions with gene strand information remained for subsequent analysis, 99% of which had maximum spliced alignment overhangs greater than 20bp.

### Euclidian distance of two frequency matrix

Supposing *f*1 and *f*2 are two 6×15 lower triangular frequency matrices, where an element in row *M* and column *k* represents the median frequency of the *k*th most dominant acceptor site of a donor site with *M* alternative acceptor sites, *k*≤*M*. The frequency matrix data may be derived from experimental RNA-seq data, or from a simulation of the Weibull distribution *W(0*.*14)*, such as in Fig 2B. The Euclidian distance of the two matrixes can be calculated from following formula:

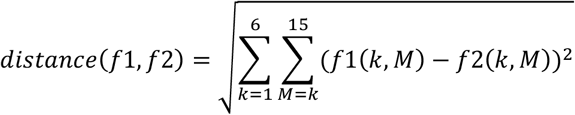

### Kullback-Leibler divergence of two distributions

Supposing P and Q are two probability density distributions — with P derived from experimental data and Q derived from simulated data of a Weibull distribution — the Kullback-Leibler divergence (KLd) between P and Q is defined by following formula:

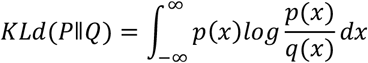

When P and Q are discrete probability distributions:

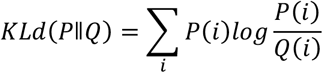

KLd represents the amount of information lost when Q is used to approximate P.

## Author Contributions

Conceptualization: J.H. Data curation: J.H. Formal analysis: J.H. Funding acquisition: D.C.D. Investigation: J.H., D.C.D. Methodology: J.H. Project administration: J.H., D.C.D. Software: J.H. Supervision: J.H. D.C.D. Validation: J.H. Visualization: J.H. Writing-original draft: J.H. D.C.D. Writing-review & edition: J.H., D.C.D.

## Supplemental Information

**Fig S1.**
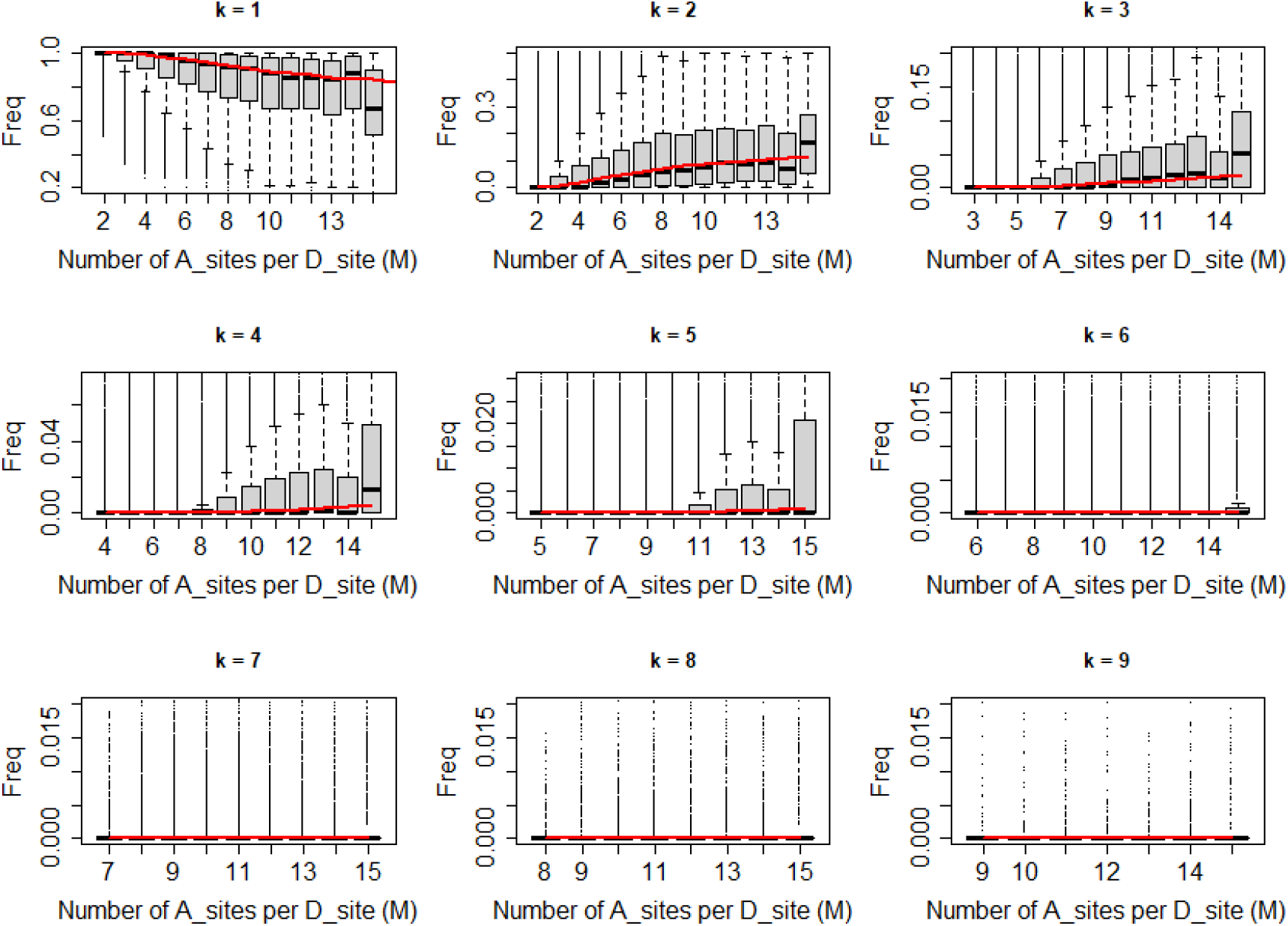
Boxplots of the distribution of frequency of alternative acceptor sites with fixed *k* and increasing *M* from RNA-seq data. *k* is the rank of alternative acceptor sites. *M* is total number of alternative acceptor sites. Red curve is median frequency calculated from *W*(0.14).

**Fig S2.**
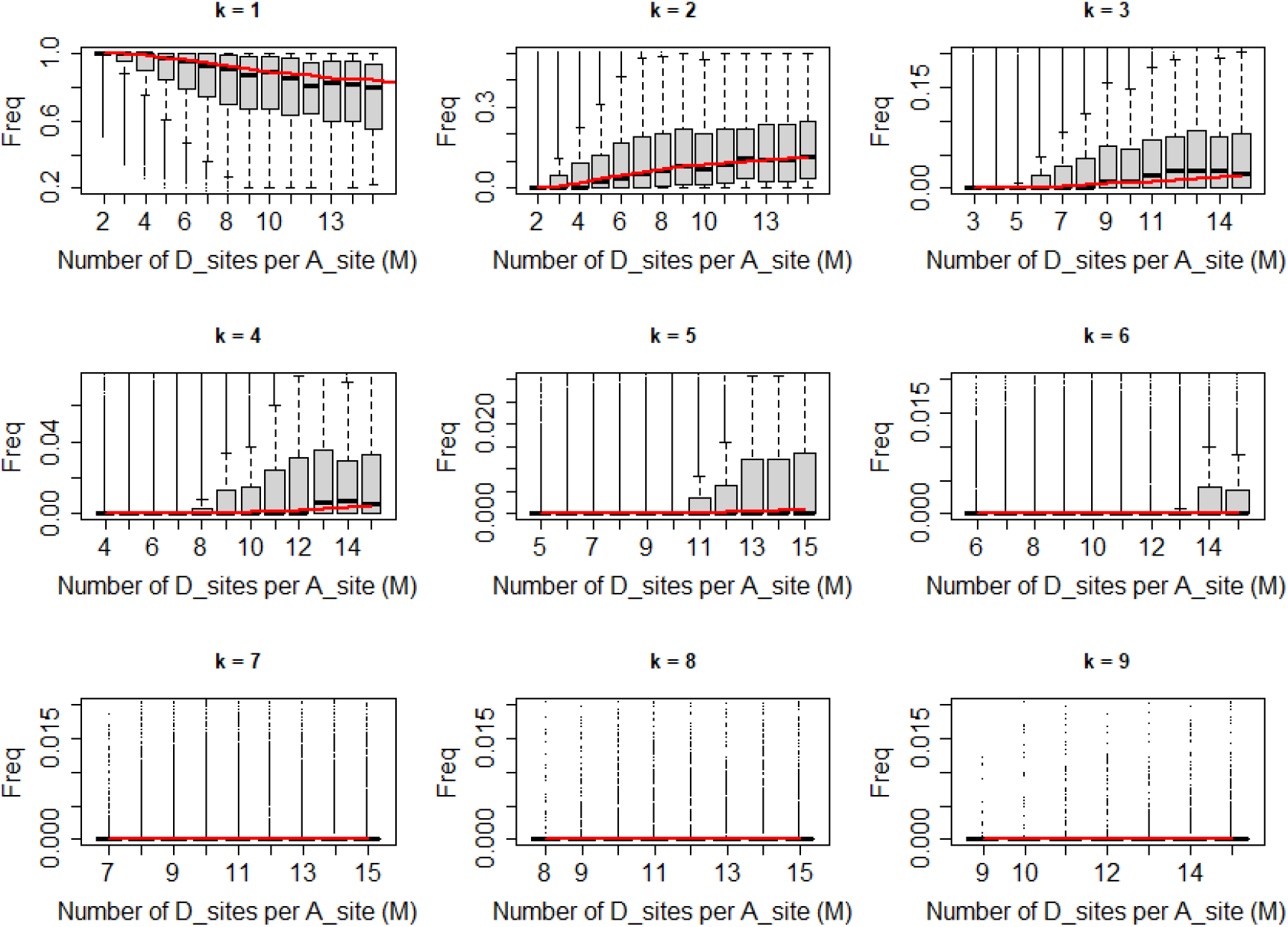
Boxplots of the distribution of frequency of alternative donor sites with fixed *k* and increasing *M* from RNA-seq data. *k* is the rank of alternative donor sites. *M* is total number of alternative donor sites. Red curve is median frequency calculated from *W*(0.14).

**Fig S3.**
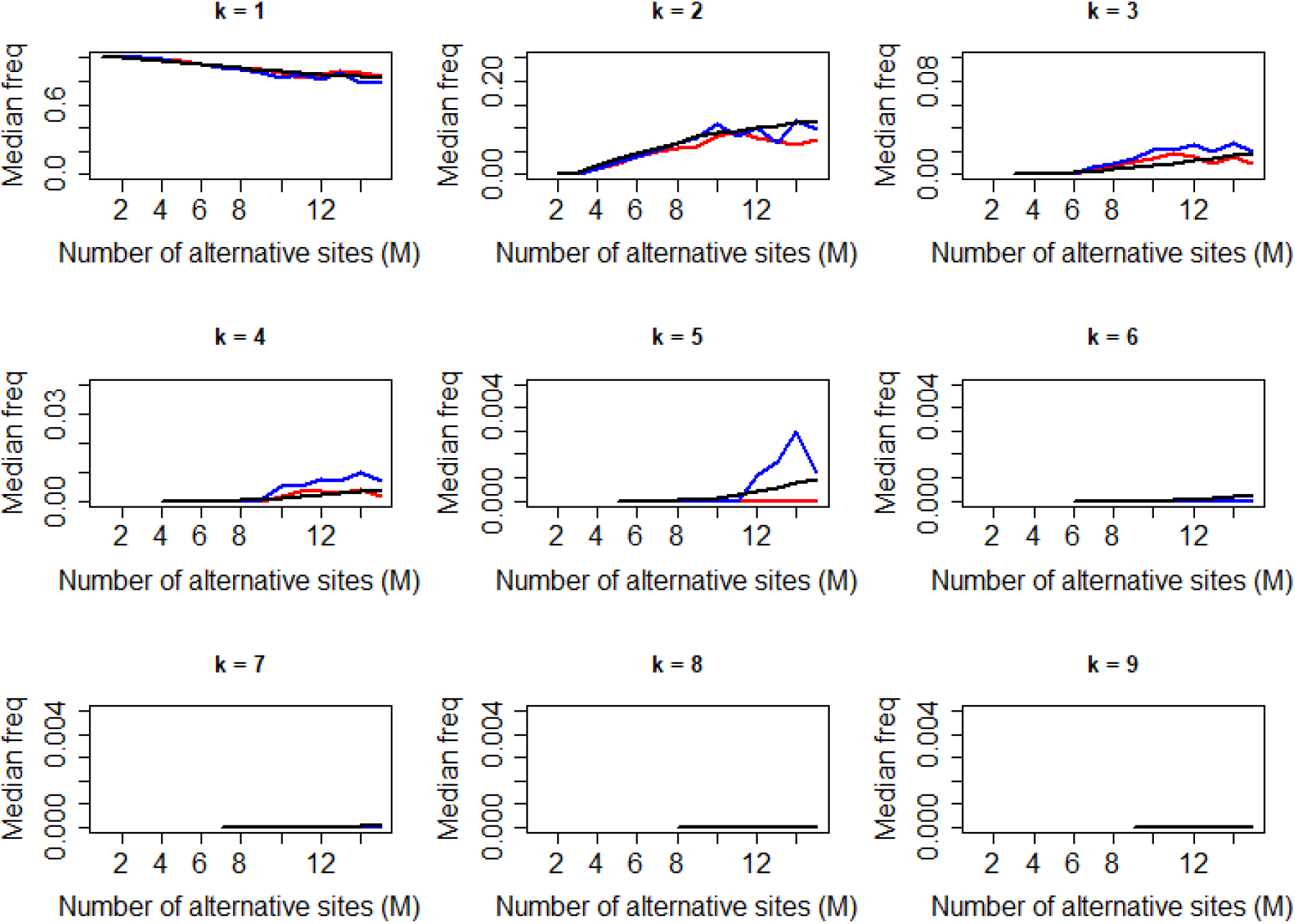
The distribution of median frequency of alternative acceptor and donor sites with fixed *k* and increasing *M* from colorectal cancer RNA-seq data (GSE50760). *k* is the rank of alternative splice sites. *M* is total number of alternative splice sites. Red line represents alternative acceptor sites. Blue line represents alternative donor sites. Black line represents simulated distribution from *W*(0.14).

**Fig S4.**
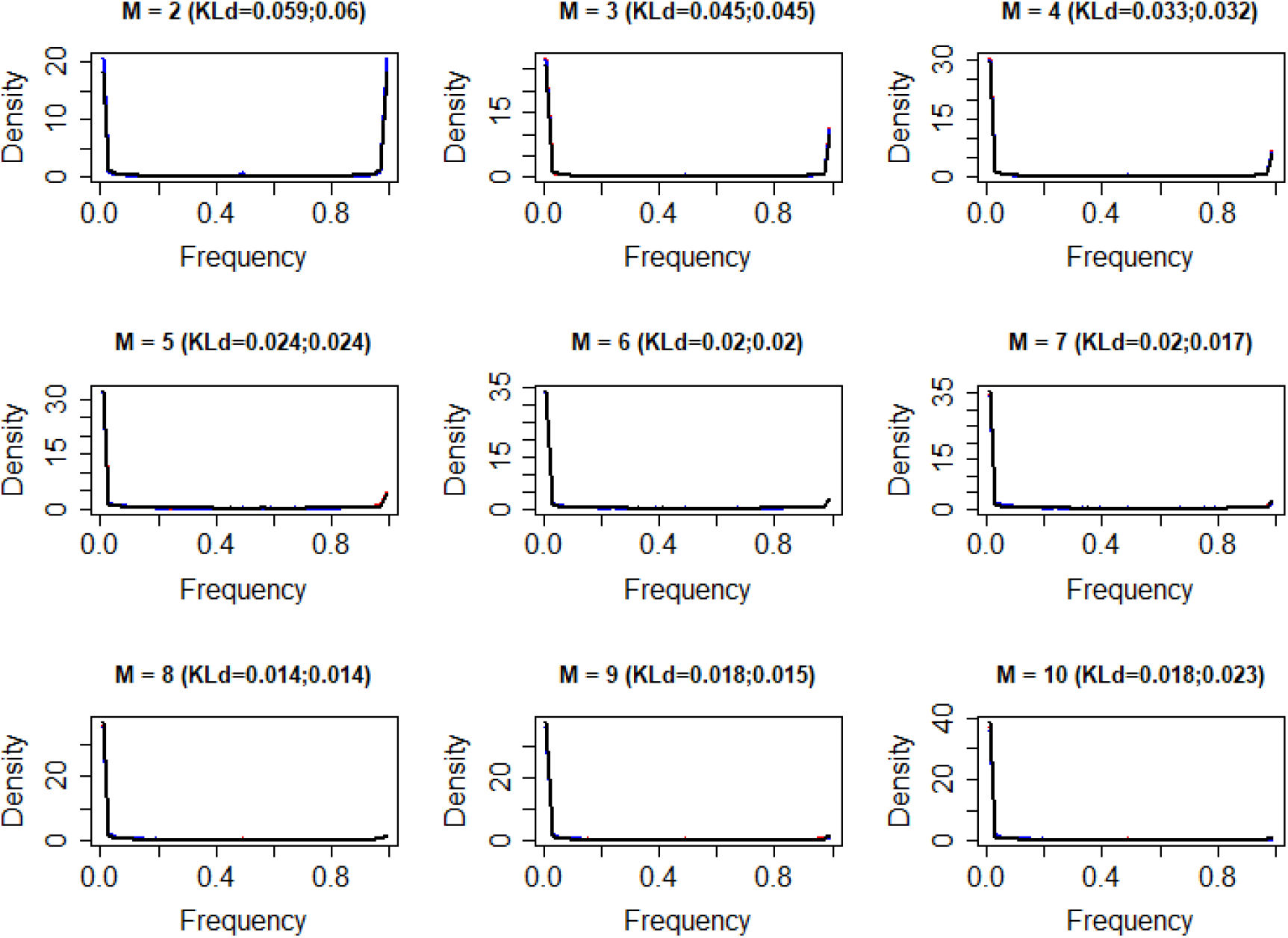
The frequency distribution of all alterative splice sites from colorectal cancer RNA-seq data (GSE50760) and simulated data for M=2:10. M is the number of alternative donor and acceptor sites. Red curves represent alternative acceptor sites, blue curves represent alternative donor sites, black curves represent simulated data from *W*(0.14). KLd is the Kullback-Leibler divergence between the two distributions. The first number in parentheses is KLd between the frequency distribution of alternative acceptor sites and simulated data from *W*(0.14). The second number in parentheses is KLd between the frequency distribution of alternative donor sites and simulated data from *W*(0.14). The three curves almost exactly overlap.

**Fig S5.**
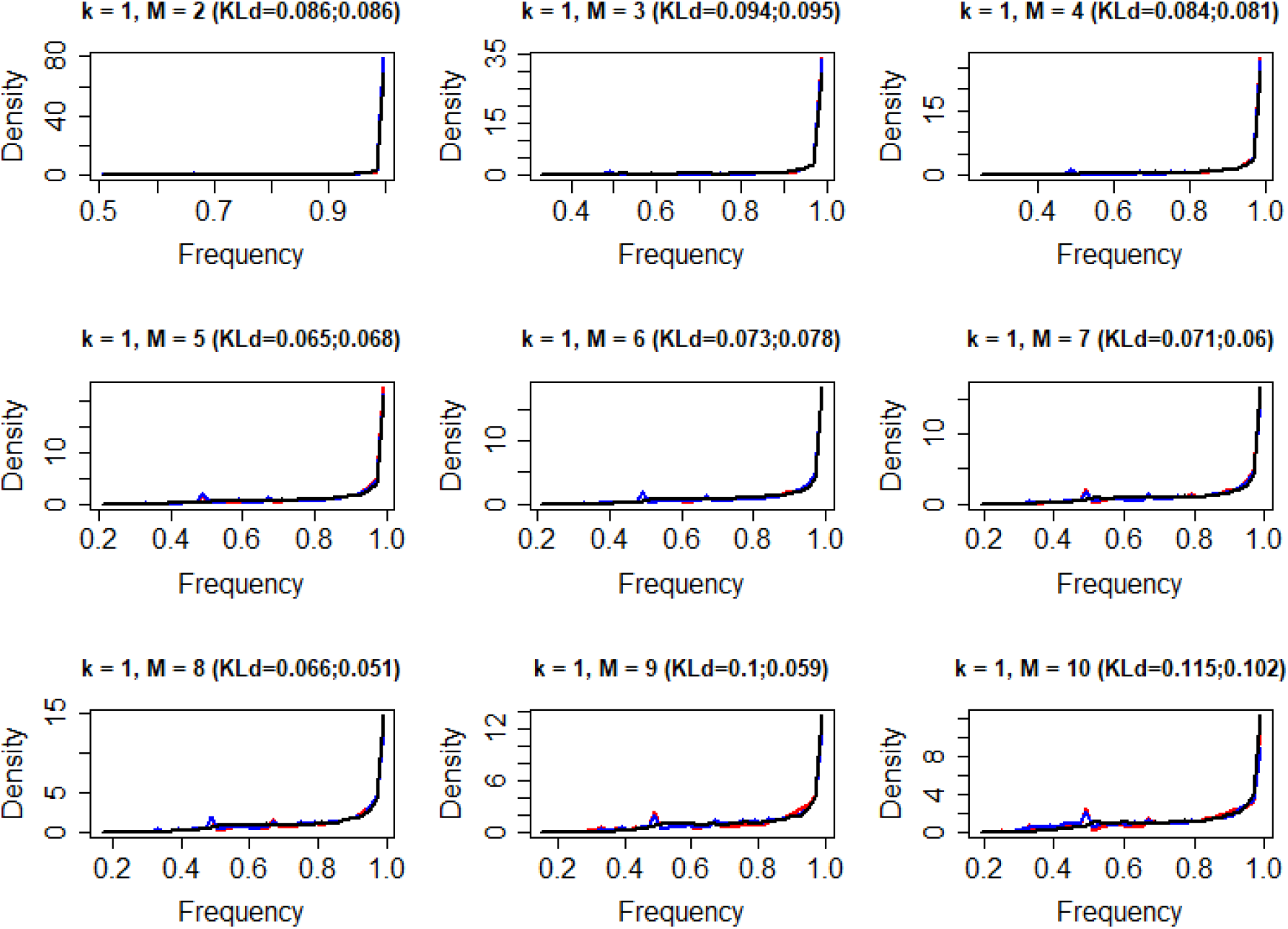
The frequency distribution of the most dominant alterative splice sites (k=1). M is the number of alternative donor and acceptor sites. The experimental data is from colorectal cancer RNA-seq data (GSE50760). Red curves represent alternative acceptor sites, blue curves represent alternative donor sites. Black curves represent simulated data from *W*(0.14). KLd is the Kullback-Leibler divergence between two distributions. The first number in parentheses is the KLd between the frequency distribution of alternative acceptor sites and simulated data from *W*(0.14). The second number in parentheses is KLd between the frequency distribution of alternative donor sites and simulated data from *W*(0.14). The three curves almost exactly overlap.

**Fig S6.**
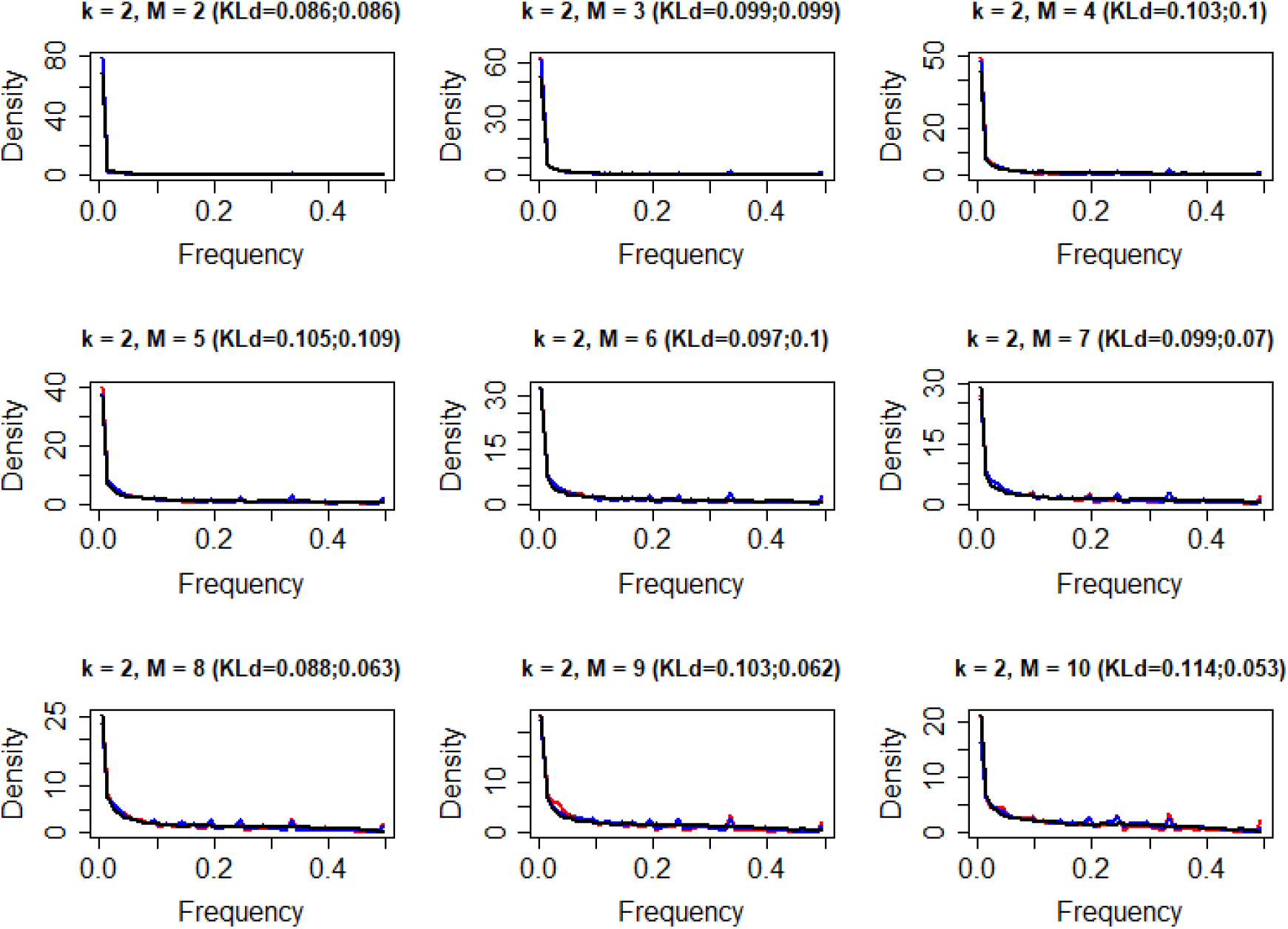
The frequency distribution of the second most dominant alterative splice sites (k=2). M is the number of alternative donor and acceptor sites. The experimental data is from colorectal cancer RNA-seq data (GSE50760). Red curves represent alternative acceptor sites, blue curves represent alternative donor sites. Black curves represent simulated data from *W*(0.14). KLd is the Kullback-Leibler divergence between two distributions. The first number in parentheses is the KLd between the frequency distribution of alternative acceptor sites and simulated data from *W*(0.14). The second number in parentheses is KLd between the frequency distribution of alternative donor sites and simulated data from *W*(0.14). The three curves almost exactly overlap.

**Fig S7.**
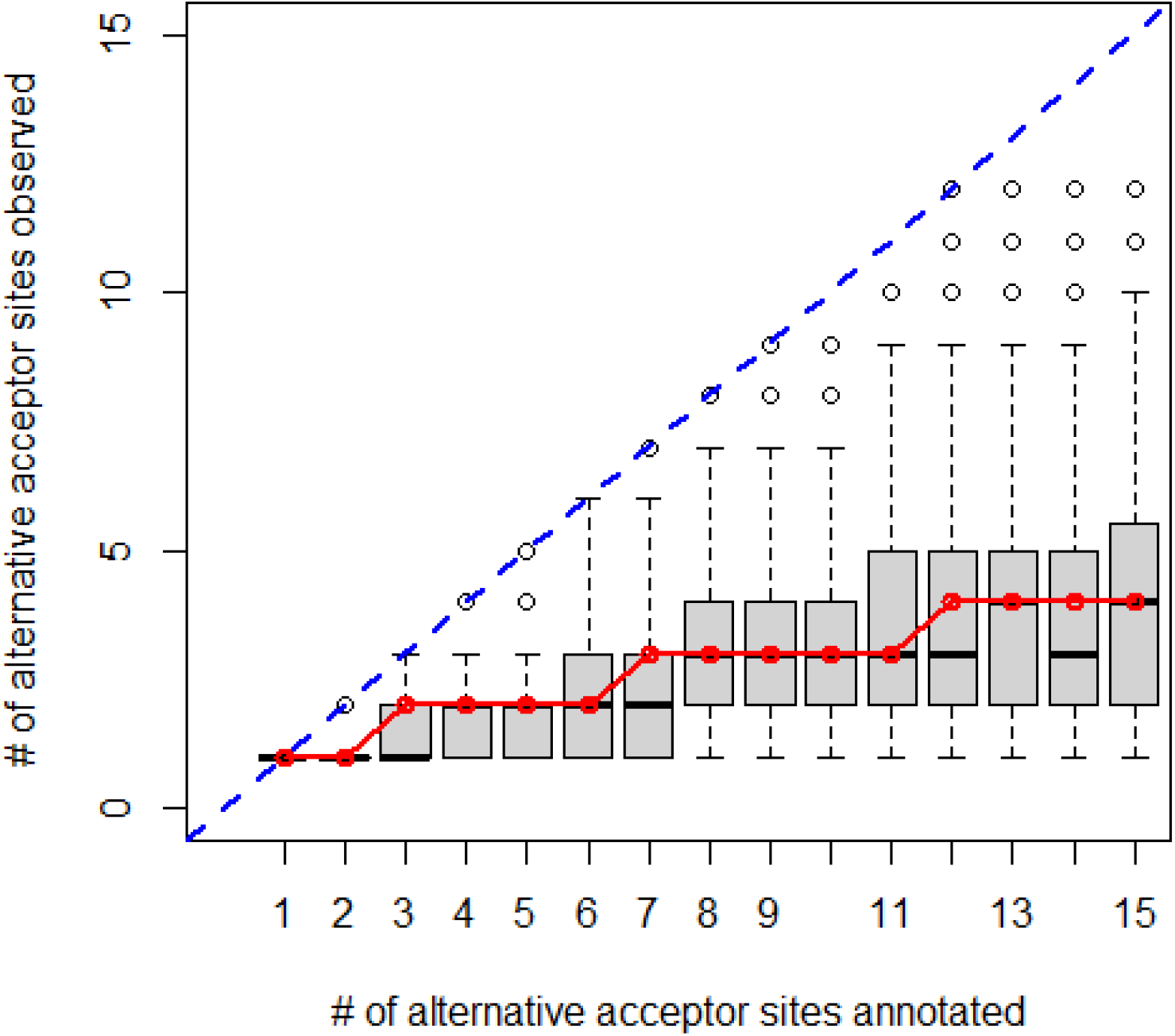
Boxplots of the distribution of the number of alternative acceptor sites observed in one sample versus those annotated. The boxplot is the observed result from our RNA-seq data. The red curve is the expected median calculated from *W*(0.14).

**Fig S8.**
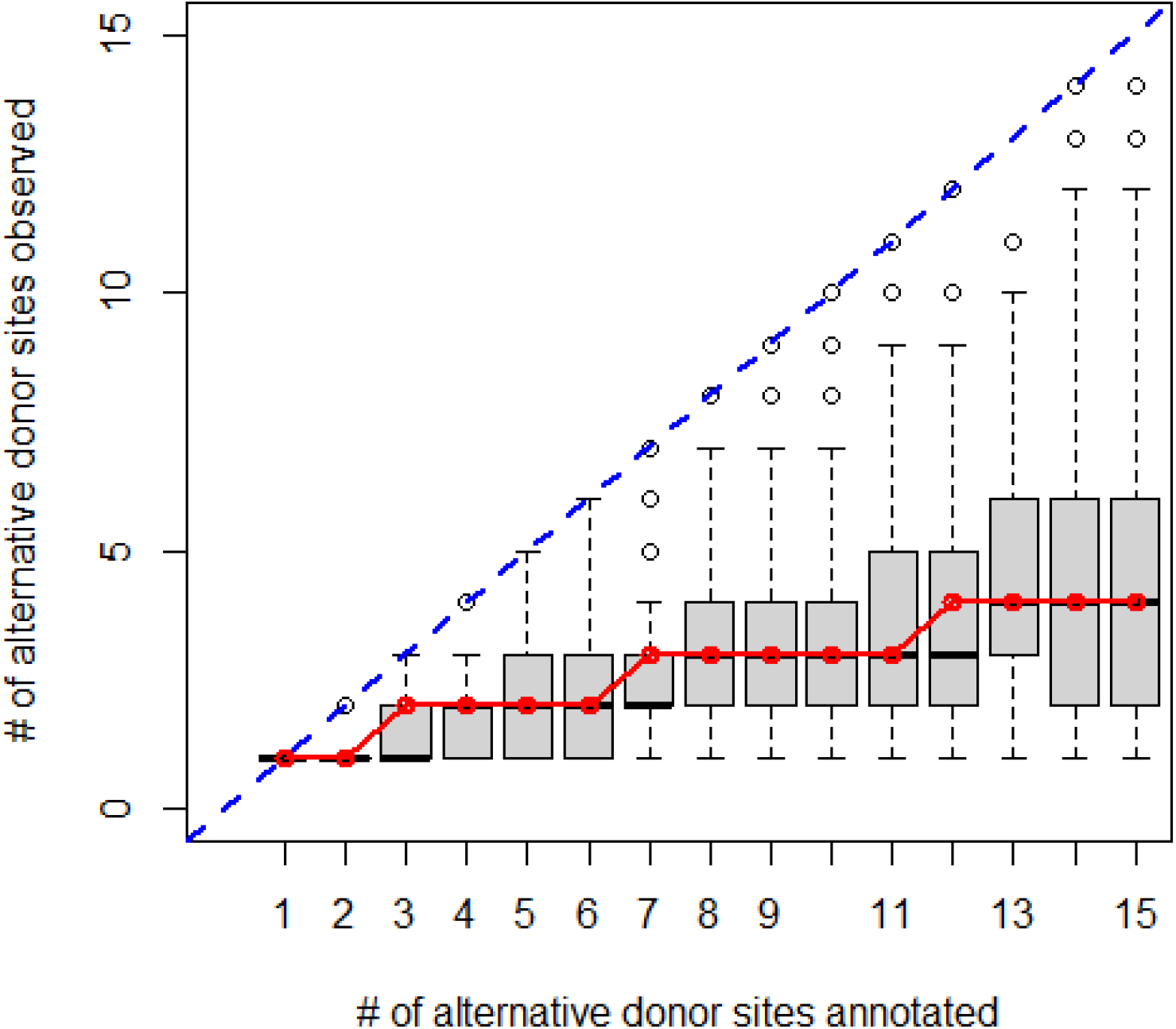
Boxplots of the distribution of the number of alternative donor sites observed in one sample versus those annotated. The boxplot is the observed result from our RNA-seq data. The red curve is the expected median calculated from *W*(0.14).

## Notes

### Competing Interest Statement

The authors have declared no competing interest.

### Summary of Updates

Corrected name error and removed line number

